# Stool Dynamics and the Developing Gut Microbiome During Infancy

**DOI:** 10.1101/2025.06.30.662361

**Authors:** Mohammed Al-Andoli, Sarah Schoch, Andjela Markovic, Christophe Mühlematter, Matthieu Beaugrand, Oskar G. Jenni, Rabia Liamlahi, Jean-Claude Walser, Dennis Nielsen, Salome Kurth

**Affiliations:** Department of Psychology, University of Fribourg, Fribourg, Switzerland; Department of Pulmonology, University Hospital Zurich, Zurich, Switzerland; Center of Competence Sleep & Health Zurich, University of Zurich, Zurich, Switzerland; Child Development Center’ University Children’s Hospital Zurich, Zurich, Switzerland; Genetic Diversity Center, ETH Zurich, Zurich, Switzerland; Department of Food Science, University of Copenhagen, Copenhagen, Denmark

## Abstract

The infant gut microbiome is a dynamic ecosystem that plays a crucial role in early development, influences immune system maturation, and overall health. Recent insights reveal that the gut microbiota undergoes changes across the 24-h day, raising the possibility that it may act as “zeitgeber”, helping to regulate the host’s sleep-wake organisation. Despite its importance, timing factors influencing microbiome composition are poorly understood, limiting its use as a health predictor. This study investigates the relationship between stool dynamics (interval since last defecation, collection time of the sample), sleep pressure (interval since last sleep), meal timing and the composition of the gut microbiome. Stool samples from 198 healthy infants, aged 3 to 31 months, were analysed to assess microbial diversity, richness evenness, and abundance. Our findings reveal that longer intervals since the last bowel movement are associated with increased microbial diversity, evenness, and richness. Stool timing is associated with shifts in microbial composition, especially in younger infants, indicating that diurnal microbial fluctuations may become more stable as infants mature. We found that longer periods of wakefulness were associated with increased microbial diversity in early infancy, although this effect appeared to diminish with age. Feeding schedules appeared to have a limited effect on the gut microbiome. Longer fasting before sampling showed no significant associations with most microbial parameters, except for a positive association with microbial richness. At the phylum level, results indicate that infant gut microbial composition is influenced by behaviour and physiology. Longer intervals between bowel movements were associated with shifts in bacterial abundance, with *Proteobacteria* decreasing and *Actinobacteria* increasing. Stool timing and meal timing also affected microbial diversity: later sampling times showed higher *Actinobacteria* levels, and longer fasting was associated with reduced *Bacteroidetes*. Sleep pressure showed a trend effect with *Firmicutes* displaying a slight decrease in infants who had been awake longer. Our findings underscore the importance of time-based factors on infant gut microbiome composition.

## 1. Introduction

The infant gut microbiome plays a key role in early development, influencing overall health by supporting immune system maturation, metabolic processes, brain development and sleep regulation [1-3]. In turn, sleep is essential for facilitating key physiological processes such as growth, immune function, and neurodevelopment across the lifespan [4]. Insufficient sleep duration disrupts these processes, negatively affecting metabolism, behaviour, and increasing the risk of various diseases [5]. Poor sleep quality in infancy and early childhood has been linked to long-term consequences, including obesity, asthma or cognitive dysfunction [6, 7].

The gut microbiome shows substantial variation between individuals during the first year of life [4, 8], influenced by factors such as feeding type, delivery method (e.g., vaginal or cesarean) and antibiotic use [9]. Diurnal factors, such as diet and sleep pattern, also influence gut microbiome dynamics [4, 8]. Despite its critical role in overall health, many determinants of microbiome development remain poorly understood. Gaining deeper insight into the complex interactions between the gut microbiome, stool dynamics, meal timing, and sleep could lead to more personalized clinical approaches in paediatric care.

The human gut microbiota is a complex ecosystem composed of bacteria, eukaryotes, archaea and viruses, including bacteriophages (phages) that target bacteria [10]. This microbial community plays essential roles in protecting against pathogens, modulating the immune system, producing vital metabolites, and nourishing gastrointestinal cells [11]. Crucially, a significant portion of the microbiota undergoes rhythmic oscillations, with up to 60% of its composition fluctuating periodically [12]. In mice, around 20% of commensal species exhibit diurnal variations, and a similar trend is observed in humans, where 10% of species show fluctuations across the day [12]. Specific microbial taxa, such as *Clostridiales*, and *Bacteroidetes* in mice, and *Parabacteroides, Lachnospira*, and *Veillonella* in humans, are known to follow these oscillations [13]. These findings suggest that the gut microbiota may play a role in the host’s internal clock, with microbiota rhythmicity becoming more pronounced with age and influenced by early-life dietary patterns [14].

Factors, such as the timing of sleep, feeding, and bowel movements are closely linked to diurnal or circadian rhythms, likely influencing both the function and composition of the gut microbiome [15, 16]. Circadian rhythm is an internal process, that governs the sleep-wake organisation over a 24 hour period and affects various physiological functions, including gastrointestinal activity [4]. The internal circadian rhythm is calibrated by external “zeitgebers”, contextual cues that support rhythms of physiological processes and thus also the behavioural rhythmisation of sleep-wake organisation [17]. In adults, disruptions to circadian rhythm have been linked to alterations in gut microbiota composition, which can negatively impact metabolic health [18, 19].

The complex relationship between the developing gut microbiome and its surrounding environment - including factors like the specific clock time of stool collection, and the frequency of bowel movement patterns - is an area of growing scientific interest [4, 16]. Early infancy represents a critical window when the gut microbiota undergoes rapid growth and development [14], and yet despite its importance for several pathways of health, the influence of stool dynamics on microbiome composition remains largely unexplored. These interactions are generally poorly understood at any age of the human lifespan, yet infants, with their frequent bowel movements, offer an excellent model for examining these dynamics. This study aims to quantify the effect of temporal dynamics on infant gut microbiome composition. Specifically, this study investigates how stool dynamics (interval since last defecation, clock time of stool collection), alongside sleep pressure and meal timing shape the composition of the gut microbiome during the most critical developmental window. We hypothesize that these temporal factors significantly shape microbial composition. Data from 198 healthy infants aged 3 to 31 months were analysed, and 504 of their stool samples served as basis to characterize these associations.

We hypothesize that early infancy features a highly responsive microbiome that is sensitive to variations in the most fundamental zeitgebers, with sleep pressure and fasting periods being key factors determining individual variability of gut microbiome composition [17]. Further, as infants mature, we expect the microbiome to stabilize, becoming progressively less susceptible to external deviations and developing stronger internal rhythmicity. Accordingly, we anticipate that the timing factors (bowel frequency, stool timing, sleep pressure, fasting), affect the microbial community structure with age-specific effects, including dynamics in the relative abundance at the phyla level (*Proteobacteria, Actinobacteria, Firmicutes, Bacteroidetes*).

## 2. Methodology

### 2.1. Participants

This study included 198 healthy infants residing in Switzerland, assessed for sleep-wake patterns and bowel movements at various ages between 3 and 31 months. The predominant assessment ages were 3 months, 6 months, and 12 months (cohorts SDEGU, SPIN, NUTR), while one cohort (SLEEPY) was assessed at varying ages within 5 to 31 months. All infants were generally healthy, born full-term via vaginal delivery and had not received antibiotics during their first three months of life. Additionally, none of the mothers reported a history of gut-related disorders such as irritable bowel syndrome (IBS), inflammatory bowel disease (IBD), or *Clostridioides difficile* infection [20]. A more detailed overview of the inclusion and exclusion criteria is available in [21].

The study was conducted in accordance with ethical guidelines, with approval granted by the Cantonal Ethics Committees, Switzerland (BASEC 2016-00730, 2019-02250). All parents or primary guardians provided written informed consent after receiving a comprehensive explanation of the study procedures, ensuring compliance with principles outlined in the Declaration of Helsinki.

### 2.2. Experimental Design

A total of 504 stool samples were collected from four independent cohorts: SDEGU (n= 389, from 160 infants, 117 at age 3 mo, 135 at age 6 mo, 137 at age 12 mo), SPIN (n=43, from 12 infants, 19 at age 3 mo, 24 at age 6 mo), NUTR (n=31, from 12 infants, 16 at age 3 mo, 15 at age 6 mo), and SLEEPY (n=41, from 14 infants, 5-31 mo, with a mean sampling age of 12.2±6.2 months, mean±SD). Parents utilized disposable pipettes and spatulas to collect stool samples from diapers, which were then stored in sterile tubes inside protective bags for temporary refrigeration. Within 72 hours, the samples were transported in cooling boxes to the laboratory, where aliquots were stored at -50°C or -80°C for subsequent analysis [20]. Data on gut microbiota composition were obtained from stool samples through 16s rRNA gene amplicon sequencing, targeting the V3 region to characterize bacterial communities. Contextual information for each stool sample was either provided by the parents or derived from a 24-hour diary they completed across several continuous days during the study [20].

Our analysis focused on four key factors that could potentially create variance in infant gut microbiome composition: two related to stool dynamics - the time since the last bowel movement (bowel frequency) and the clock time of stool passage – and two additional factors: the time elapsed since the last sleep (sleep pressure) and time since the last meal (meal timing), both derived from the parent-completed 24-hour diary entries.

### 2.3. Characterization of gut microbiota

DNA was extracted from approximately 200 mg of stool using the PowerSoil® kit, with minor modifications. After heat treatment (65°C for 10 minutes, followed by 95°C for 10 minutes), the samples were bead-beaten to lyse bacterial cells. The DNA extraction steps adhered to the manufacturer’s protocol. 16S rRNA gene amplicons targeting the V3 region were generated using specific primers compatible with the Nextera Index Kit® (Illumina). Further details on amplification, barcoding, and sequencing are provided in [1, 4]. A subsample of this data (SDEGU cohort) has been described in [20].

A comprehensive data processing and analysis pipeline was employed to identify zero-radius Operational Taxonomic Units (zOTUs) using UNOISE [22]. Denoising and taxonomic assignment of amplicons were performed using USEARCH::UNOISE3 and USEARCH::Sintax with the SILVA SSU (v138) database. To improve data quality, samples with ‘time since last stool’ greater than 48 hours were excluded. Data normalization was performed through rarefaction to standardize sequencing depth across samples. Negative controls identified a potential contaminant (zOTU), but this had minimal effect on the final results. Core bacteria were defined as those present in at least 20% of samples with a minimum relative abundance of 1% [14].

Three key metrics were used to assess the gut microbiome: diversity, evenness, and richness. Diversity (Shannon index) reflects the variety of species in a sample, with higher values representing a more diverse community. Evenness reflects how uniformly individual species are distributed, with higher evenness indicating a more balanced community. Richness (number of taxa) measures the total number of distinct species [23]. The relative abundance of zOTUs was also calculated and analyzed in relation to the four time-based factors (time since last stool, stool timing, sleep pressure, time since last meal). Abundance was determined by counting zOTUs that exceeded a relative abundance threshold of 0.01, identifying the key contributors to overall microbial diversity in each sample. Additionally, the study examined the relative abundance of bacterial phyla—*Proteobacteria, Actinobacteria, Firmicutes*, and *Bacteroidetes*—focusing on the effects of time-related factors, across the age groups.

### 2.4. Statistical analysis

Linear regression analysis was conducted to assess relationships between microbial parameters and stool dynamics. Regression models estimated the association of independent variables, i.e., time since the last bowel movement and the clock time of stool sampling, with microbial metrics, including diversity, evenness, and richness. The beta coefficient for each predictor variable was derived from the model to quantify the strength and direction of these associations. Age was included as covariates in the models to control for potential confounding effects. For repeated measures, linear mixed-effects models were employed, with random intercepts to account for within-subject variability. Correlation analysis was utilized to further explore the strength of these relationships. Analysis of Variance (ANOVA) was used to validate the findings from correlation analysis and to determine differences in microbial parameters based on variations in independent variables. Both linear regression and Pearson correlation were used to investigate the associations between microbial parameters and stool dynamics across three age group. In addition, these statistical tests were used to examine the effects of stool dynamics on the composition of bacterial phyla.

## 3. Results

### 3.1. Time since an infant’s last bowel movement affects gut microbiome composition

First, we examined the association between the time since the last bowel movement and the composition of the infant gut microbiota. Samples were grouped into 3-hour intervals based on the time since the last stool (ranging from 1 to 48 hours, with samples outside this range excluded, i.e. 20 samples) to reduce noise and variability in the data. For example, samples collected within 1 to 3 hours after a bowel movement were grouped into the 3-hour time block, while those collected 4 to 6 hours afterward were represented in the 6-hour block. This method ensured a balanced representation of samples across time intervals.

The summary of microbial diversity, richness, and evenness across different time intervals since the last stool indicates that the mean Shannon diversity increases as the time interval progresses, peaking at 45 hours (mean = 2.67), with variability in standard deviation (SD) across intervals (ranging from 0.27 to 1.51). The richness values show a similar upward trend, with the highest mean richness observed at 42 hours (mean = 487.50), followed by gradual fluctuations, while the SD for richness is consistently higher than that for Shannon diversity. The mean evenness ranges from 0.38 to 0.53, with values generally increasing as the time intervals grow. The sample sizes vary across intervals, with the largest sample size at 24 hours (99 samples) and smaller sizes at extreme intervals (e.g., 2 samples at 42 and 45 hours).

Within each time block, data of samples were averaged for key microbial parameters, including diversity, evenness, and richness (Figs. 1 and 2). As shown in Fig. 1, variations in time since the last stool related to microbial parameters, specifically, we observed an increase in diversity, evenness, and richness across the first 12 hours, followed by a slight decrease between approximately 12-30 hours, and a subsequent increase after approximately 30-36 hours without stool passage, yet the later with increasing variance between individual time blocks.

**Figure 1.**
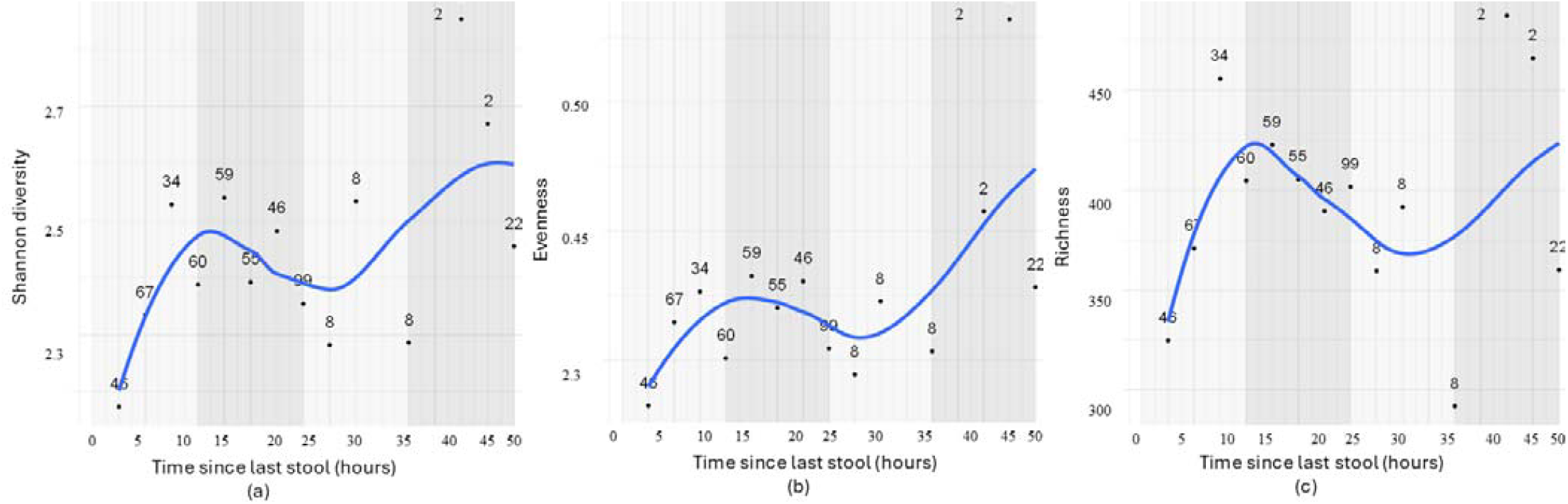
Relationship between time since the last bowel movement and gut microbial diversity (a), evenness (b), and richness (c) in infants (n=504 samples: n= 152 at 3 months, 174 at 6 months, 137 at 12 months, and n=41 at 5-31 months). Samples were grouped into 3-hour intervals to adjust for uneven representation of sample numbers across time points. For each interval, averages of microbial parameters (diversity, evenness, and richness) were calculated. The displayed numbers indicate the number of samples included in each interval. The blue line represents the actual changes in microbiome parameters as time since the last stool changes, while the numbers in plots represent the number of samples per interval.

**Figure 2.**
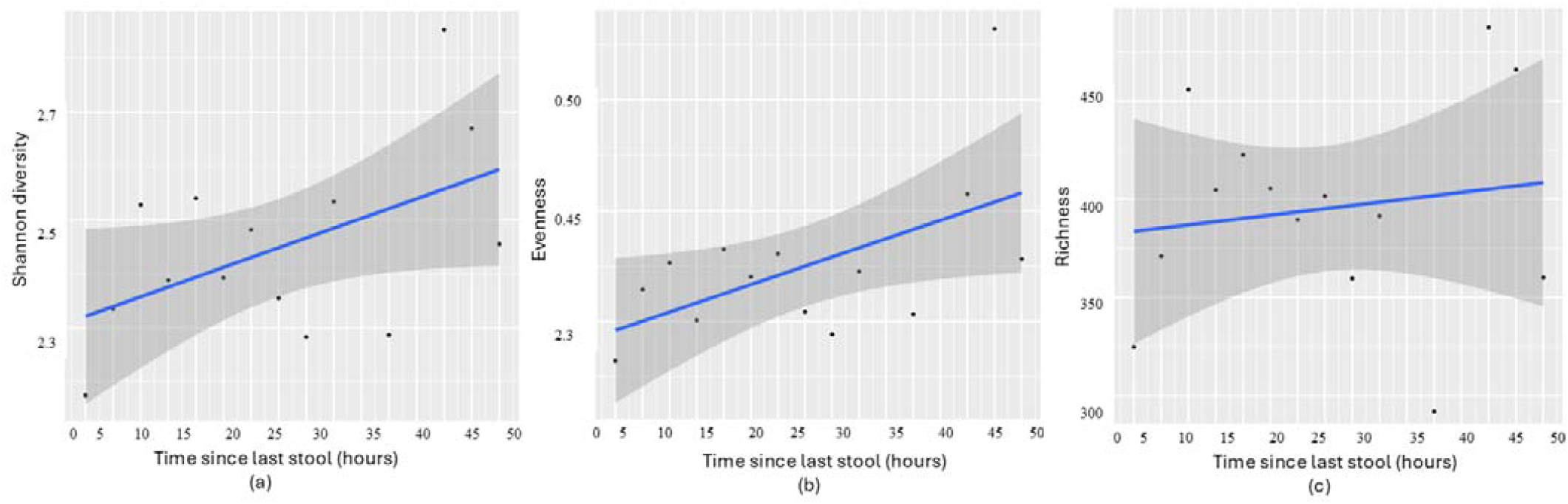
Linear regression analysis examining the relationship between time since the last stool and (a) Shannon diversity (*p* = 0.056, β = 0.018), (b) evenness (*p* = 0.038, β = 0.004), and (c) richness (*p* = 0.60, β = 1.656) in infants. Samples were grouped into 3-hour intervals, consistent with Figure 1.

Linear regression analysis revealed that longer intervals since the last bowel movement were associated with increased microbial diversity (Shannon *p* = 0.056 and β= 0.018) and evenness (*p* = 0.038, Beta β=0.004), suggesting a more diverse and balanced microbial community over time (Fig. 2). However, microbial richness did not show a significant association (*p* = 0.605, β= 1.656), indicating that the total number of different species remains stable, suggesting the presence of a core set of species that persists regardless of the time elapsed since the last bowel movement. In alignment, correlation coefficients show strong positive relationships between time since the last stool and both microbial diversity (r = 0.51, *p* = 0.056) and evenness (r = 0.56, *p* = 0.038), and there is no effect for richness (r = 0.15, *p* = 0.605).

To further examine the relationship between stool frequency and microbial parameters, a one-way Analysis of Variance (ANOVA) was conducted. The results were consistent with the findings from the regression analysis. Time since the last stool was significantly associated with Shannon diversity (F = 4.23, *p* = 0.056, and evenness (F = 5.43, *p* = 0.038; 3-hour intervals), indicating that longer intervals between bowel movements are linked to increased microbial diversity and a more even distribution of species. Again, no association was observed for richness (F = 0.29, *p* = 0.601), suggesting that while time since last stool explains variability of microbial diversity and evenness, it does not explain the dynamics in the overall number of microbial taxa present.

Next, LMM account with participant (infant) included as a random effect to account for repeated measures, and age treated as a fixed covariate to adjust for potential confounding. The analysis was used on all samples (without interval grouping) to capture variance between infants across continuous time points. This model accounts for repeated measures within the same infants across different age groups, following an approach similar to MaAsLin2 for handling longitudinal data. Regression coefficients (coef) and p-values (*p*) were computed. The analysis revealed that longer intervals since the last bowel movement were associated with increased microbial diversity (Shannon: coef = 0.065, *p* = 0.025), evenness (coef = 0.053, *p* = 0.03), and richness (coef = 0.072, *p* = 0.009), suggesting that microbial communities become more diverse and balanced over time (Fig. 2).

Results from grouped samples (Figures 1 and 2) and ungrouped samples (Figure 3) confirmed significant associations between microbial diversity and evenness with time since the last stool. Richness was only significant in the ungrouped analysis, likely due to preserved individual variability.

**Figure 3.**
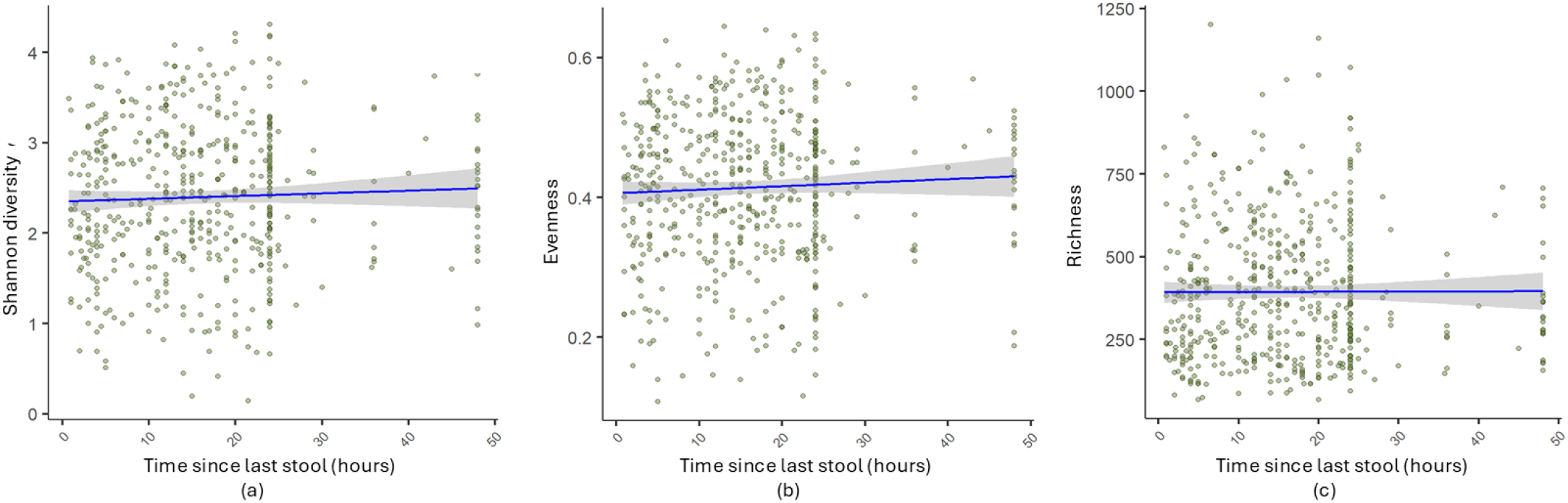
Mixed-effects model analysis of (a) Shannon diversity (*p* = 0.025, coeff = 0.065), (b) evenness (*p* = 0.031, coeff = 0.53), and (c) richness (*p* = 0.090, coeff = 0.071) in relation to time since the last stool in infants. All samples were analyzed individually without grouping into time intervals.

### 3.2. Infants stool timing affects gut microbial diversity and evenness

We next investigated the association between the timing of bowel movements (i.e. the clock time when the stool sample was obtained) and microbial parameters. Similar, to before samples within close time frames were grouped into one-hour intervals, and the corresponding gut microbial parameters were averaged (Figs. 4 and 5). The trajectories indicate, from a descriptive perspective, that variations in stool timing throughout the 24-hour-day - with data obtained within the range 3 am to 11 pm – relate to variance in microbial parameters. Specifically, diversity and evenness increased across the second half of the night from approximately 3 am to 10 am, followed by a period of stability across the daytime until approximately 11 pm. Meanwhile, richness showed a similar pattern first - increasing approximately from 3 am to 10 am, stabilizing between 10 am and 7 pm - but then starting to decrease towards midnight. Overall, these patterns reveal the notable observation that infant gut microbial parameters exhibit the greatest fluctuation during nighttime hours.

**Figure 4.**
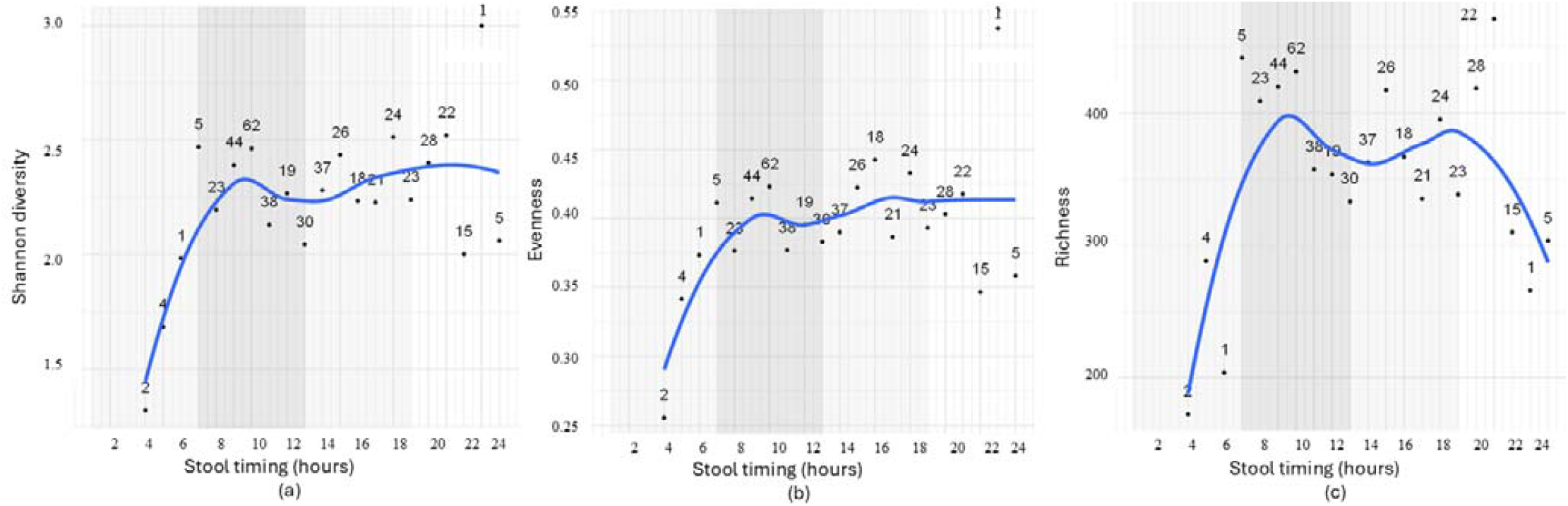
The association between stool timing and gut microbial diversity (a), evenness (b), and richness(c) in infants. The infant samples are grouped into one-hour intervals, and all samples in the same interval was averaged in term of gut microbiome parameters.

**Figure 5.**
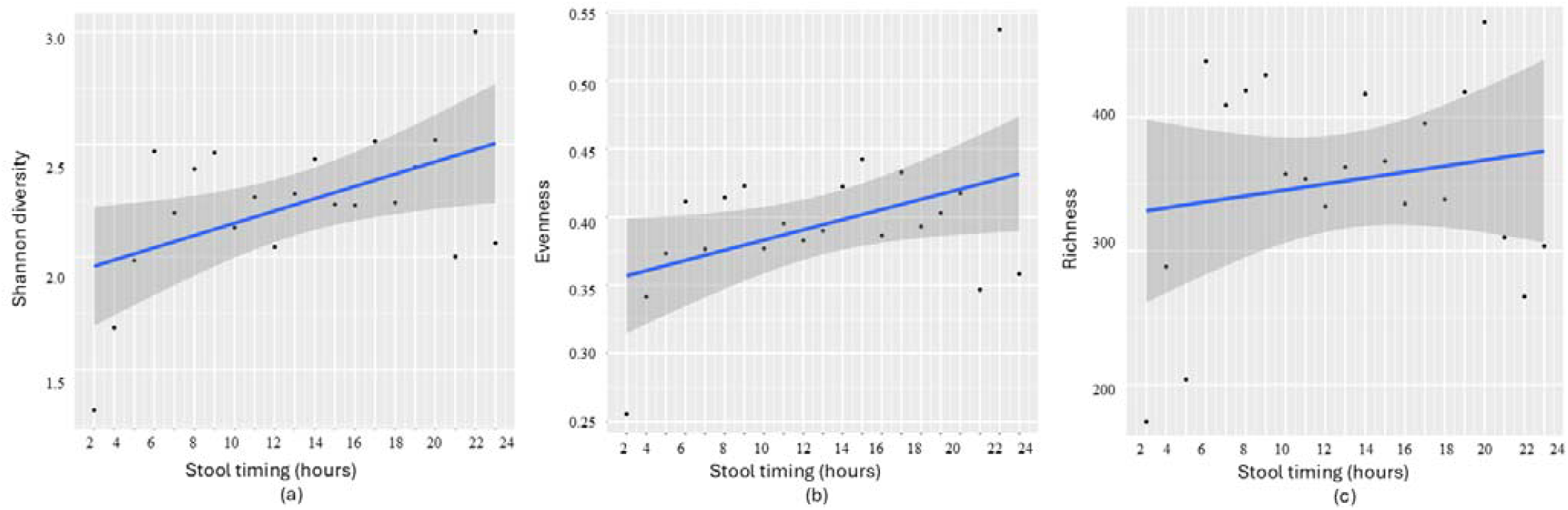
Linear regression analysis of (a) Shannon diversity (*p* = 0.056, β = 0.018), (b) evenness (*p* = 0.007, β = 0.003), and (c) richness (*p* = 0.53, β = 1.76) in relation to infant stool timing (1-24 hours). Samples were grouped into one-hour intervals.

The summary of microbial diversity, richness, and evenness across 1-hour stool timing intervals indicates that Shannon diversity generally increases with time, with some fluctuations, peaking at 22 hours (mean = 3.00, based on a single sample). Richness shows variability across intervals, with a high mean richness observed at 6 hours (441.60) and 20 hours (443.81). Evenness is relatively consistent across intervals, ranging from 0.26 to 0.54. The standard deviation (SD) highlights variability within each interval, and sample sizes vary widely, with some intervals having very few samples (e.g., 1 sample at 5 and 22 hours)

Linear regression analysis of one-hour interval samples confirmed the dynamic patterns by demonstrating a significant increase in diversity (*p* = 0.045, β= 0.024) and evenness (*p* = 0.07, β= 0.003) with later stool timing, as assessed within the timeframe 3 am to 11 pm. Richness, however, did not show a significant relationship with stool timing (*p* = 0.53, β= 1.76). Correlation analysis revealed coefficients r = 0.50 (*p* = 0.021) for diversity, r = 0.45 (*p* =0.043) for evenness, and r = 0.17 (*p* =0.434) for richness. ANOVA was conducted to further explore the relationship between stool timing and gut microbiome parameters. The analysis confirmed that stool timing affects microbial parameters, with Shannon diversity (F= 6.383, p= 0.020), and evenness (F=4.708, p= 0.042). However, no significant effect was observed for richness (F= 0.637, *p* = 0.435), indicating that stool timing does not affect the overall number of microbial species present.

LMM was applied to all samples individually (without interval grouping). No significant associations were found between stool timing and gut microbiome parameters, except for a weak negative association with richness (coef = - 0.041, *p* = 0.09). Increased variability and age adjustment likely masked trends observed in grouped data.

### 3.3. Age-related differences in stool patterns and gut microbiome parameters based on stool dynamics across three infant age groups

In the next step, we examined how age may influence the relationship between stool timing and gut microbiome parameters, by focusing on longitudinal changes in the three cohorts with repeated within-subject sampling (3, 6, and 12 months, i.e. cohorts SDEGU n=387, SPIN n=43, and NUTR n=31; Fig. 6). The infant samples are grouped into three-hours intervals in time since last stool and one-hour intervals in stool timing variable. We examined the association of stool timing and microbiome composition separately for each age group with liner regression analysis (Table 1).

**Table 1.** Linear regression between stool timing parameters and microbial parameters represented for each age group (p-values *p;* Pearson correlation *r*)

|  | (a) Time since last stool |  |  |  |  |  | (b) Stool timing |  |  |  |  |  |
| --- | --- | --- | --- | --- | --- | --- | --- | --- | --- | --- | --- | --- |
|  | 3 Months |  | 6 Months |  | 12 Months |  | 3 Months |  | 6 Months |  | 12 Months |  |
| | $p$ | $r$ | $p$ | $r$ | $p$ | $r$ | $p$ | $r$ | $p$ | $r$ | $p$ | $r$ |
| Diversity | **0.021** | **0.66** | **0.033** | **0.59** | 0.554 | 0.21 | **0.019** | **0.52** | 0.224 | 0.27 | 0.179 | -0.34 |
| Richness | 0.223 | 0.46 | 0.368 | 0.2993 | 0.229 | 0.42 | 0.079 | 0.42 | 0.609 | 0.11 | 0.067 | -0.45 |
| Evenness | **0.035** | **0.57** | 0.156 | 0.3992 | 0.653 | 0.16 | 0.084 | 0.38 | 0.454 | 0.14 | 0.228 | -0.31 |

**Figure 6:**
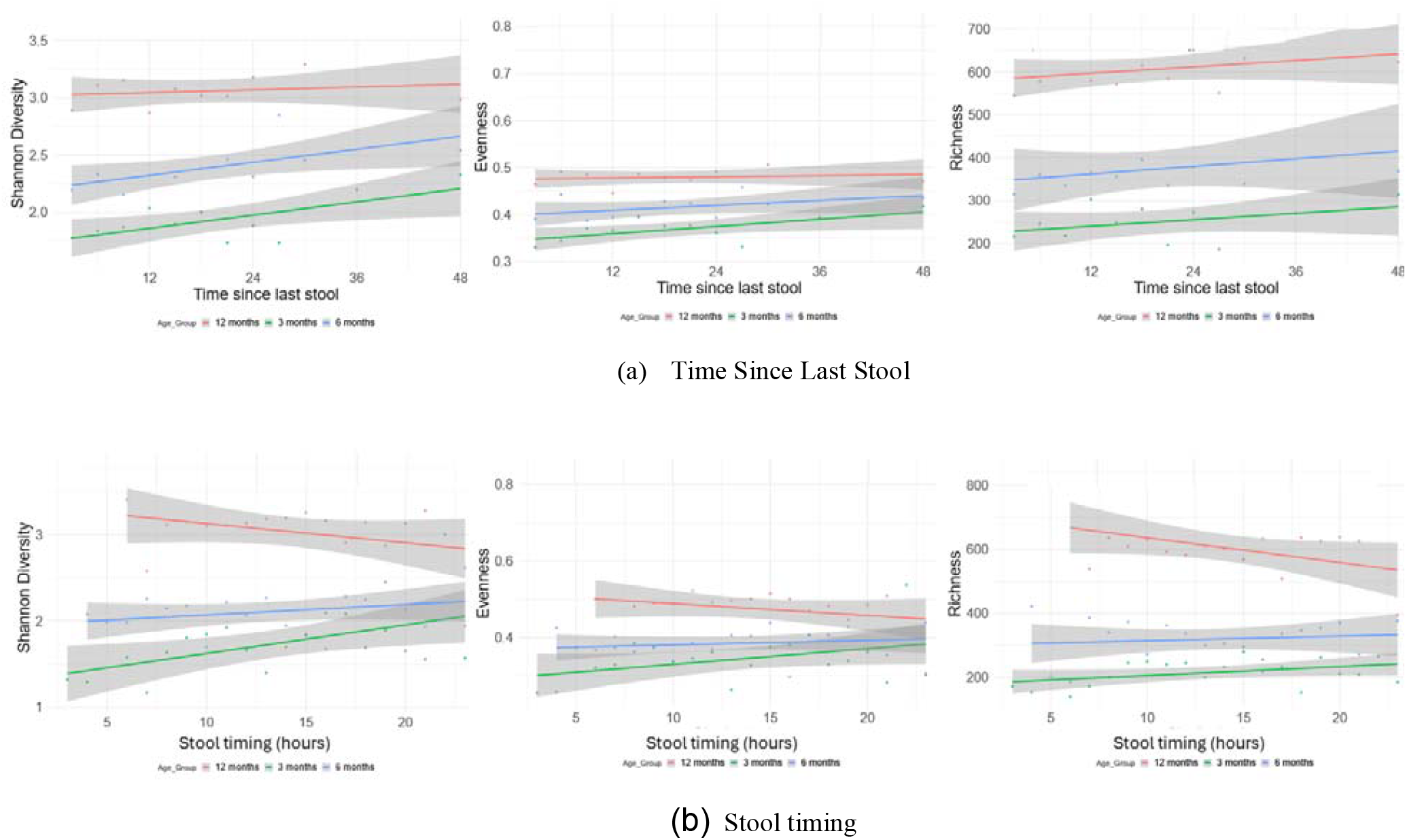
Stool dynamics and gut microbiome parameters (diversity, evenness, and richness) across three timepoints (3, 6, and 12 months). The infant samples are grouped into three-hours intervals in (a) time since last stool and one-hour intervals in (b) stool timing variable. The associations between (a) time since last stool and (b) stool timing with microbiome parameters are summarized by p-values reported in Table 1.

Time passed since last bowel movement was a significant factor explaining microbial variance at age 3 months, showing a positive linkage with microbial diversity and evenness. At 6 months, this effect prevailed with microbial diversity, yet by 12 months, no significant associations were observed. These data reveal strong effects of bowel frequency on microbial composition at the youngest age, and a gradual vanishing across the first year of life.

Similar relationships were observed for stool timing (clock stool): At 3 months, later stool timing was linked to increased microbial diversity, and trends prevailed for evenness and richness. At 6 months, effects disappeared, and by 12 months, notably, richness revealed a negative association reaching trend-level significance.

Next, the analysis was applied to infant samples individually (without interval grouping) across three infant age groups. Significant associations were observed at 3 months between gut microbiome parameters and time since the last stool (Figures 7a-c), as well as weak links between stool timing and gut microbiome parameters (Figures 7d and e). At 3 months (Figures 7a-c), time since the last stool was linked to increased microbial diversity, with trends also observed for richness and evenness. Stool timing at 3 months was weakly linked to richness and evenness but showed no association with diversity. At 6 and 12 months, no significant associations were found with any gut microbiome parameters. The results from ungrouped infant samples based on time intervals support the findings from grouped samples discussed in Figure 6a and 6b.

**Figure 7.**
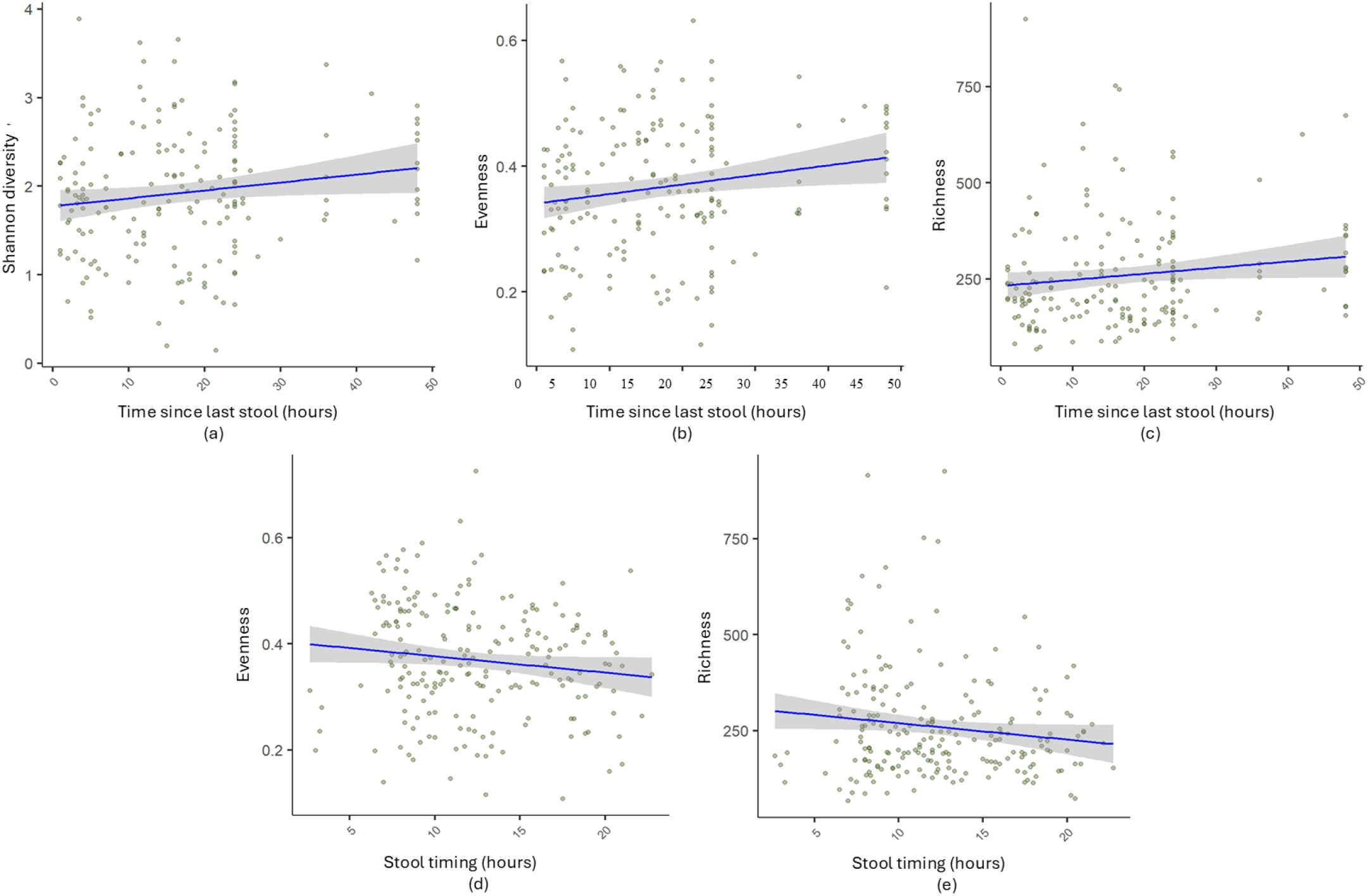
Stool dynamics and gut microbiome parameters at 3 months, using all samples individually without interval grouping. Panels (a–c) show associations between time since last stool and gut microbial diversity (*p* = 0.081, coeff = 0.09), evenness (*p* = 0.022, coeff = 0.079), and richness (*p* = 0.016, coeff = 0.12), respectively. Panels (d–e) display associations between stool timing and gut microbial evenness (*p* = 0.012, coeff = –0.05) and richness (*p* = 0.14, coeff = –0.071).

### 3.4. Sleep pressure and fasting periods determine infant gut microbiome parameters

We then investigated whether variations in infant gut microbiome composition are associated with the duration of prior wakefulness (sleep pressure) and fasting (time since last meal). This analysis was conducted using 504 samples across all cohorts. Samples were grouped into 1-hour intervals, and the gut microbial parameters (diversity, evenness, richness) were averaged within each 1-hour interval.

Samples within up to 6 hours after the last sleep episode were included (exclusion of 9 outlier samples). Sleep pressure was significantly associated with all microbial dynamics: Longer wakefulness before sampling was linked to increased diversity (p=0.0002, r=0.54), richness (p= 0.001, r= 0.48) and evenness (p=0.023, r=0.35), as shown in Table 2. Detailed time trajectories suggest an immediate and rapid increase in diversity, a delayed rise in richness, beginning approximately 3 hours later, and a gradual increase in evenness (Fig. 8a-c).

**Table 2.** Linear regression (p-values *p*, correlation coefficients *r*) of the associations among sleep pressure (time since last sleep) and time since last meal with microbial composition parameters.

| Variables | Sleep pressure |  | Time since last meal |  |
| --- | --- | --- | --- | --- |
| | $p$ | $r$ | $p$ | $r$ |
| Diversity | **0.0002** | **0.54** | 0.280 | 0.20 |
| Richness | **0.001** | **0.48** | **0.009** | **0.47** |
| Evenness | **0.023** | **0.35** | 0.738 | 0.06 |

**Figure 8.**
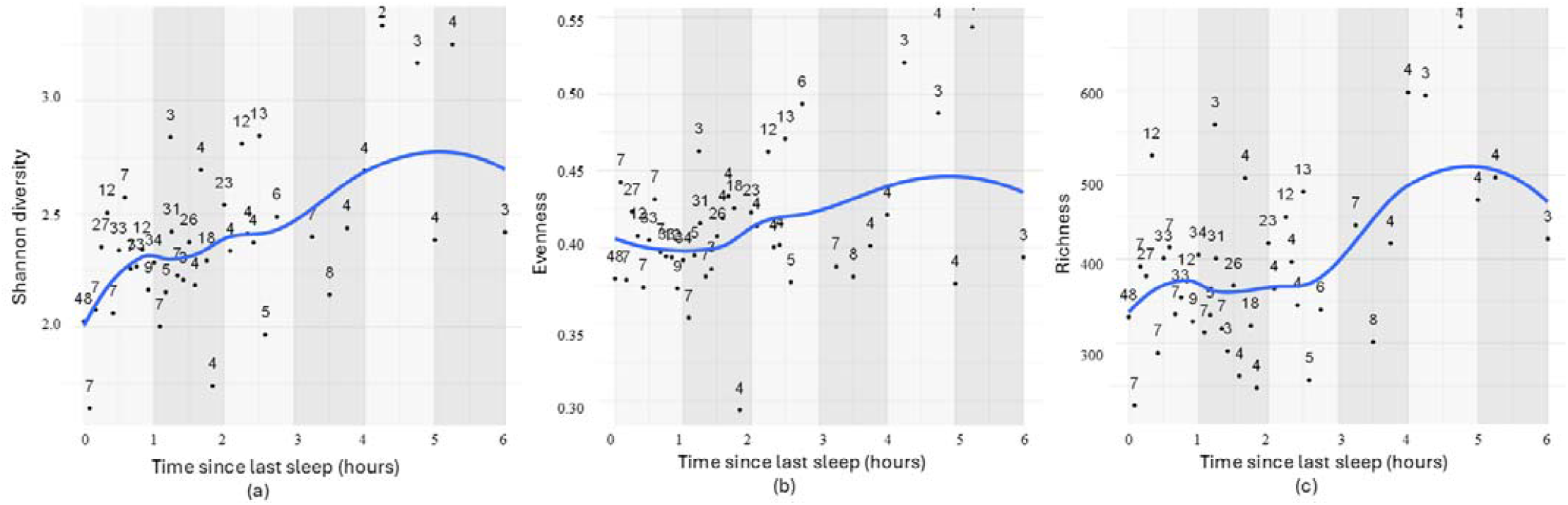
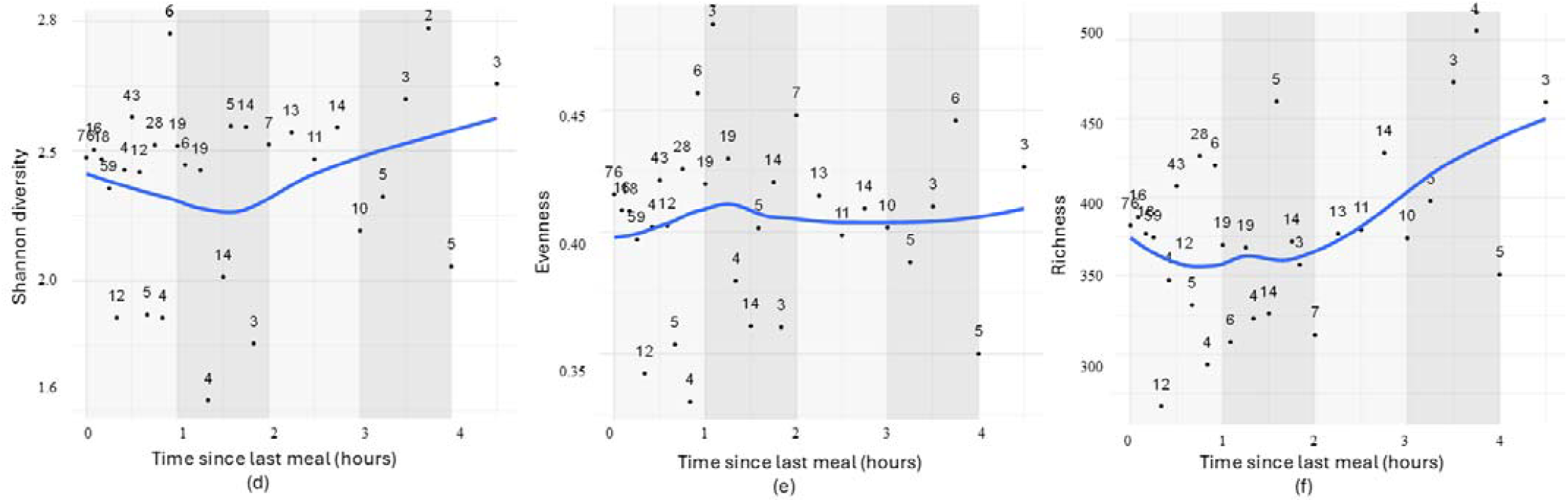
Relationship between sleep pressure (time since last sleep) and time since last meal with gut microbial diversity (a), evenness (b), and richness (c), and time since last meal with gut microbial diversity (d), evenness (e), and richness (f). All samples within a 15-min time interval were grouped and averaged for gut microbial parameters (diversity, evenness, richness).

Most samples were collected within the first two hours after the last meal. With longer fasting, only bacterial richness displayed a statistically significant increase (Table 2). In line with this, detailed time trajectories suggest minimal dynamics in diversity and evenness in the immediate aftermath of a meal or fasting. However, the increase in richness becomes more apparent after 2 hours and beyond (Figs. 8d-f).

To determine whether the effect of sleep pressure or time since last meal on microbial parameters is driven by age, we analysed the three time points separately (i.e., 3, 6, and 12 months from cohorts SDEGU, SPIN, NUTR), using a total of 461 stool samples (SDEGU n=387, SPIN n=43, NUTR n=31).

Indeed, age-related effects were observed in among sleep pressure and microbial diversity: We observed associations of sleep pressure with microbial diversity at 3 months (trend-level) and 6 months, but not 12 months. Neither effects for richness nor for evenness were observed at any age (Table 3). In other words, data indicate that long wakeful periods relate to a more diverse internal microbial landscape in young infants until 6 months but not beyond this age.

**Table 3.**
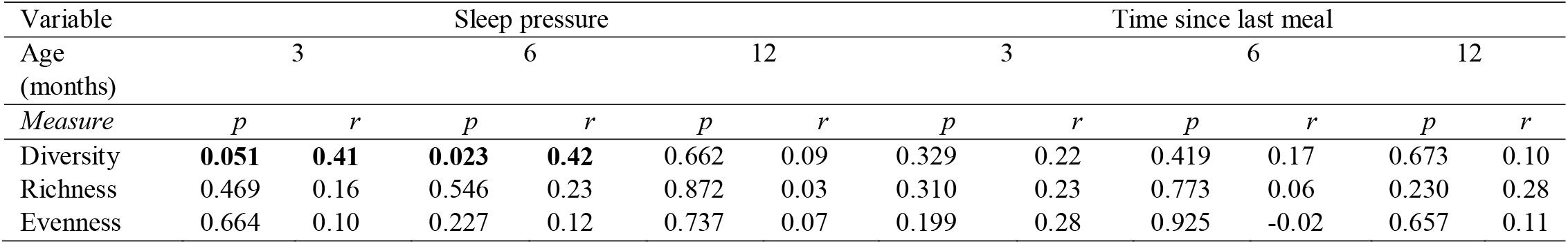
Linear regression p-values (*p*) and correlation (*r*): Association between time-based factors (time since last sleep, and time since last meal) and infant microbiome composition.

Analyses of time since last meal unravelled that the above noted effect of fasting time on microbial richness in the total sample with combined age groups disappeared when analysing by age group, identifying age as a confounder in the effect fasting may elicit on microbial richness.

Next, LMM analysis revealed no significant associations between sleep pressure and microbiome parameters. Shannon Diversity showed a weak association (β = 0.03, *p* = 0.13), while richness (β = -3.08, *p* = 0.365) and evenness (β = 0.001, *p* = 0.497) were non-significant. Similarly, “time since last meal” exhibited no significant associations, with weak trends for Shannon Diversity (β = -0.02, *p* = 0.504), richness (β = -6.73, *p* = 0.358), and evenness (β = <-0.00, *p* = 0.749).

ANOVA revealed significant associations between age and sleep pressure on diversity (F = 167), evenness (F = 76.25), and richness (F = 272), as well as between age and time since last meal on diversity (F = 25), evenness (F = 76), and richness (F = 88.4) (p < 0.0001 for all), highlighting substantial developmental changes in the gut microbiome between 3, 6, and 12 months.

We extended our analysis to evaluate associations between gut microbiota diversity and timing variables at the individual sample level. While the interval-based grouping revealed several associations, these largely diminished when analyzed ungrouped. Across all ages, weak associations were observed between sleep pressure and both Shannon diversity (*p* = 0.076) and evenness (*p* = 0.068). At 3 months, sleep pressure showed a significant association with evenness (*p* = 0.038) and a marginal association with Shannon diversity (*p* = 0.08). No associations were found at 6 months, while a negative significant link between sleep pressure and microbial diversity re-emerged at 12 months (*p* = 0.034). Regarding feeding history, only non-significant negative associations with Shannon diversity (*p* = 0.06) and evenness (*p* = 0.062) were detected at 12 months. These findings suggest that while grouping by timing intervals may reveal general trends, inter-individual variability can attenuate these associations at the single-sample level.

### 3.5. Associations between time-based factors and zOTUs relative abundance and bacterial phyla

We then refined the analysis by incorporating the relative abundance of zOTUs to achieve a more precise understanding of underpinnings of microbial diversity. Abundance was measured by counting active zOTUs (i.e., that exceeded a relative threshold of 0.01), focusing on the key contributors in each sample. Positive relationships between zOTU relative abundance and time since last stool (*p* = 0.023, *r* = 0.62), stool sampling time (*p* = 0.014, *r* =0.53), and sleep pressure (*p* = 0.008, r=0.36) were observed, while time since the last meal showed no significant effect (*p* = 0.45, *r* = 0.1; Fig. 9).

**Figure 9.**
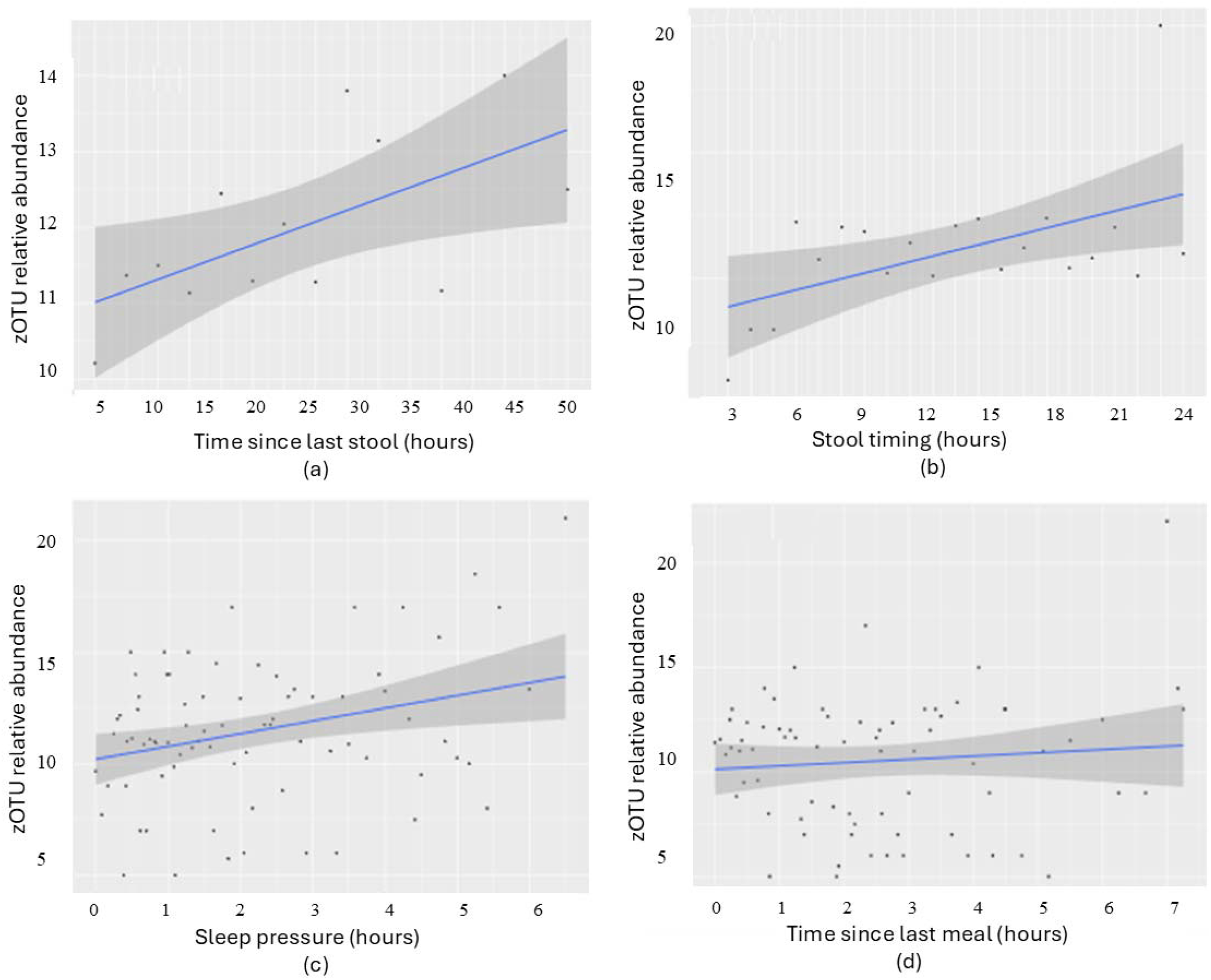
Linear regression analysis of infant group samples with relative abundances >0.01, examining associations with temporal factors: (a) Time since last stool (*p*=0.023), (b) Stool clock time (*p* = 0.014), (c) Sleep pressure (*p* = 0.008), and (d) Time since last meal (*p* = 0.425). Samples (n=504, including all ages and cohorts) were grouped into time intervals: 3-hour intervals for (a), 2-hour intervals for (b), and 1-hour intervals for (c) and (d)).

To validate these findings at the individual sample level, we extended the analysis and confirmed a robust association between abundance and time since last stool (*p* = 0.004). A marginal relationship was also observed with sleep pressure (*p* = 0.07), further supporting the influence of physiological and behavioral timing cues on microbial composition, while stool timing and meal timing remained non-significant.

### 3.6. Timing effects on major gut bacterial phyla in infants

Next, we extended the analysis to the relative abundance of the major bacterial phyla - *Proteobacteria, Actinobacteria, Firmicutes*, and *Bacteroidetes*. The analysis included all samples across ages (n=504; 152 at 3 months, 174 at 6 months, 137 at 12 months, and 41 between 5 and 31 months).

The time since the last stool related negatively with *Proteobacteria* abundance (*p* = 0.027, r = -0.15), indicating lower *Proteobacteria* levels with lower stool frequency. Conversely, Actinobacteria showed a modest positive correlation with time since last stool (*p* = 0.056, r = 0.13), indicating higher levels with lower stool frequency. Among clock time, sleep pressure and time since last meal – no relationship was detected at the phyla level (Figure 10 and Table 4).

**Table 4.**
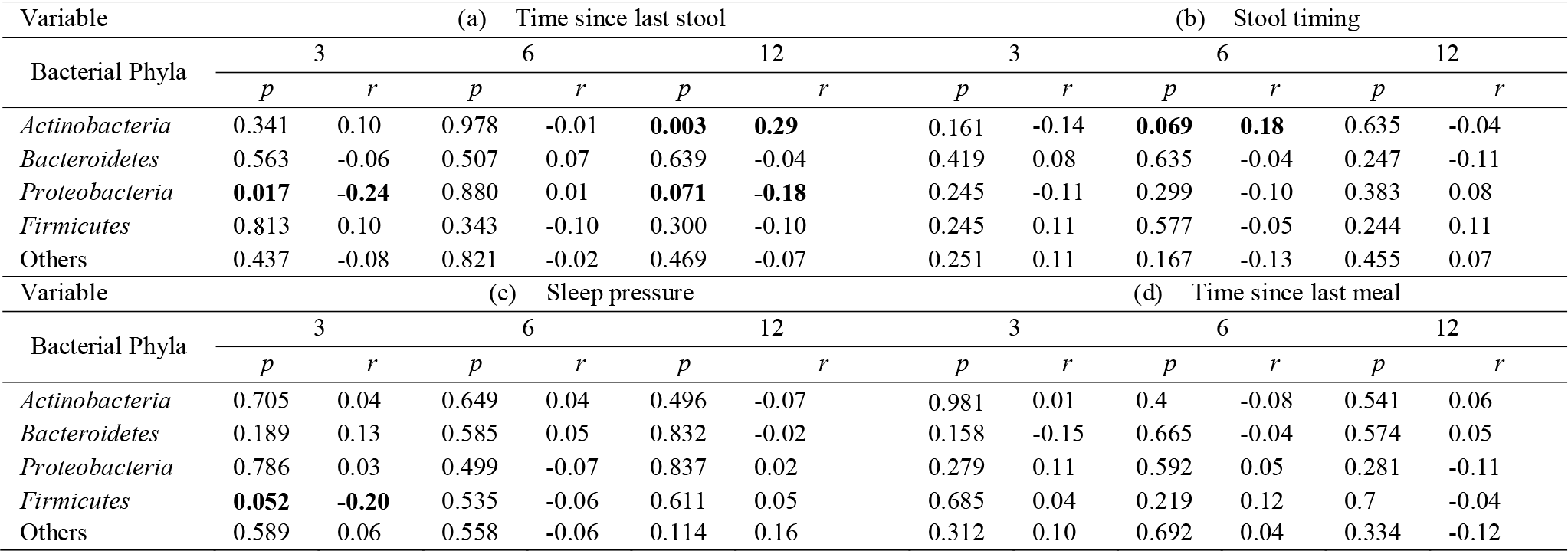
Linear Regression p-values (*p*) and correlation (*r*): Relative abundance of bacterial phyla in the infant gut by time-related factors across three age groups (3, 6, and 12 months).

**Figure 10.**
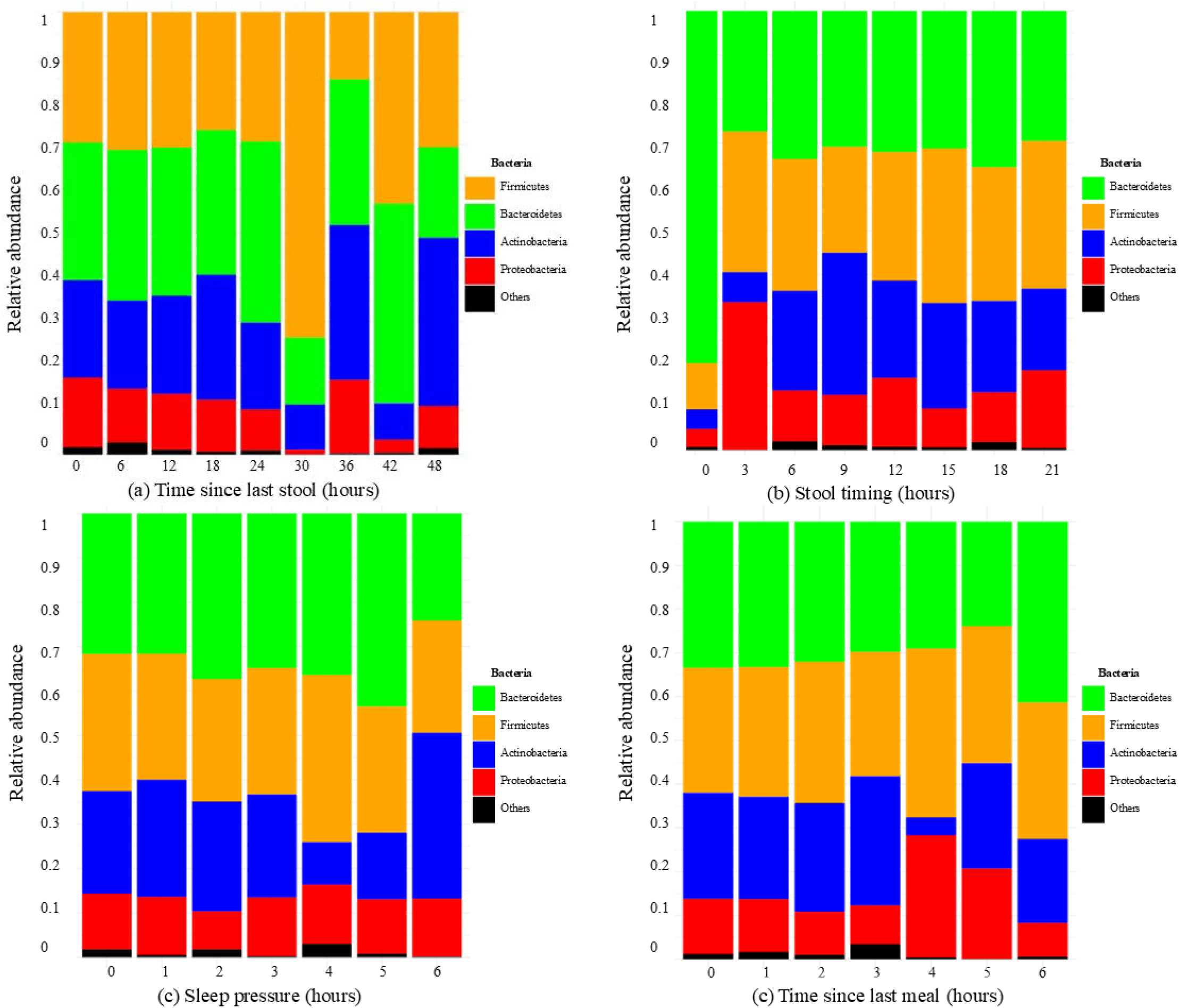
Relative abundance of major bacterial phyla in the infant gut by (a) time since last stool, (b) stool timing, (c) sleep pressure, and (d) time since last meal. Samples are grouped into 6, 3, 1, and 1-hour intervals for l, respectively, to adjust for unevenly represented hours along the x-axis.

To further differentiate temporal effects on microbiome maturation, we conducted analyses in relation to age timepoints (3, 6, 12 months)

Primarily bowel frequency was a relevant determinant: lower stool frequency was linked to decreased *Proteobacteria* at 3 and 12 months (the later at trend-level; Table 4). At 12 months, lower stool frequency was linked to increased *Actinobacteria*. Second, the timing of stool (clock stool) revealed some link to bacterial phyla, shown in a trend-level association with *Actinobacteria* at age 6 months only. Third, sleep pressure recorded an association with bacterial phyla relative abundance, concerning a decrease in *Firmicutes* with longer time wake at age 3 months. Finally, time since last meal did not exhibit any association with bacterial phyla relative abundance.

Overall, the analysis shows that effects of temporal factors on the gut microbiome vary considerably across infancy. This highlight age-specific transitions in the shaping of gut microbiota.

To control for multiple comparisons and reduce the likelihood of false positives, we applied False Discovery Rate (FDR) correction to our analyses of associations between the four time-based factors in Table 4, and the results were reported in Table 5. After FDR correction, only *Actinobacteria* at age 12 months remained statistically significant (*p* = 0.045, Table 4a), suggesting a robust association where less frequent stool is linked to increased *Actinobacteria* levels.

**Table 5.** Original p-values (*p*) and False Discovery Rate corrected p-values (*FDR*) for bacterial phyla in the infant gut by time-related factors across three age groups (3, 6, and 12 months).

| Variable | (a) Time since last stool |  |  |  |  |  | (b) Stool timing |  |  |  |  |  |
| --- | --- | --- | --- | --- | --- | --- | --- | --- | --- | --- | --- | --- |
|  | 3 |  | 6 |  | 12 |  | 3 |  | 6 |  | 12 |  |
|  | *p* | *FDR* | *p* | *FDR* | *p* | *FDR* | *p* | *FDR* | *p* | *FDR* | *p* | *FDR* |
| Bacterial Phyla |  |  |  |  |  |  |  |  |  |  |  |  |
| *Actinobacteria* | 0.341 | 0.844 | 0.978 | 0.978 | **0.003** | **0.045** | 0.161 | 0.470 | **0.069** | 0.470 | 0.635 | 0.635 |
| *Bacteroidetes* | 0.563 | 0.844 | 0.507 | 0.844 | 0.639 | 0.871 | 0.419 | 0.568 | 0.635 | 0.635 | 0.247 | 0.470 |
| *Proteobacteria* | **0.017** | 0.127 | 0.880 | 0.942 | **0.071** | 0.355 | 0.245 | 0.470 | 0.299 | 0.498 | 0.383 | 0.568 |
| *Firmicutes* | 0.813 | 0.942 | 0.343 | 0.844 | 0.300 | 0.844 | 0.245 | 0.470 | 0.577 | 0.635 | 0.244 | 0.470 |
| Others | 0.437 | 0.844 | 0.821 | 0.942 | 0.469 | 0.844 | 0.251 | 0.470 | 0.167 | 0.470 | 0.455 | 0.568 |
| Variable | (c) Sleep pressure |  |  |  |  |  | (d) Time since last meal |  |  |  |  |  |
|  | 3 |  | 6 |  | 12 |  | 3 |  | 6 |  | 12 |  |
|  | *p* | *FDR* | *p* | *FDR* | *p* | *FDR* | *p* | *FDR* | *p* | *FDR* | *p* | *FDR* |
| Bacterial Phyla |  |  |  |  |  |  |  |  |  |  |  |  |
| *Actinobacteria* | 0.705 | 0.837 | 0.649 | 0.837 | 0.496 | 0.837 | 0.981 | 0.981 | 0.4 | 0.750 | 0.541 | 0.750 |
| *Bacteroidetes* | 0.189 | 0.837 | 0.585 | 0.837 | 0.832 | 0.837 | 0.158 | 0.750 | 0.665 | 0.750 | 0.574 | 0.750 |
| *Proteobacteria* | 0.786 | 0.837 | 0.499 | 0.837 | 0.837 | 0.837 | 0.279 | 0.750 | 0.592 | 0.750 | 0.281 | 0.750 |
| *Firmicutes* | **0.052** | 0.780 | 0.535 | 0.837 | 0.611 | 0.837 | 0.685 | 0.750 | 0.219 | 0.750 | 0.7 | 0.750 |
| Others | 0.589 | 0.837 | 0.558 | 0.837 | 0.114 | 0.837 | 0.312 | 0.750 | 0.692 | 0.750 | 0.334 | 0.750 |

## 4. Discussion

Understanding the interplay between time-based factors and gut microbiota composition in early childhood is critical for advancing our knowledge of microbial development, which this study sought to examine. We analyzed 509 stool samples from 198 healthy infants, examining their immediate context, including time since last stool, sleep pressure, meal history, and clock time of stooling. Our results unveil five new insights: 1) existence of a linkage between a gut microbial composition and bowel movement time, 2) evidence age-specific transition in how gut microbiome parameters are affected by temporal factors, 3) strong associations between sleep pressure and gut microbiome composition, 4) yet no linkage with fasting periods, 5) associations of bowel movement frequency with zOTU abundance and bacterial phyla.

This comprehensive study unravelled an interplay between bowel movement frequency and various microbial parameters. Specifically, longer intervals since the last stool were associated with increased microbial diversity and evenness. These results align with existing research highlighting the infant gut microbiome’s sensitivity to bowel movement frequency [24, 25]. The observed rise in microbial diversity and evenness may reflect nutrient depletion prior defecation, indicating that the gut microbiome requires time to recover and re-establish diversity [26]. We further report that zOTU relative abundance is significantly associated by temporal factors, reflecting the dynamic nature of the gut microbiome in response to physiological changes. A positive association between time since the last stool and zOTU abundance suggests a recovery period post-defecation, during which the gut environment stabilizes, promoting microbial colonization [27]. Interestingly, the findings highlight that association between time since the last stool and gut diversity and evenness is high at younger infant ages and diminishes as infants mature. These findings suggest that the gut microbiome’s responsiveness to time-related stool factors evolves with age, highlighting a dynamic pattern of infant gut microbial development [28] .

We observed an effect of stool timing (clock time) on microbial composition. This may reflect the build-up of the enteric diurnal rhythm in the infant gut microbiome, which has been suggested based on bacterial communities in formula- and breast-fed infants [14, 29]. A recent study [14] compared bacterial communities in formula- and breast-fed infants, clock-time based assessments revealed this rhythm, yet without accounting for the timing of stool samples in relation to other potential zeitgebers, which may not fully capture drivers of microbial rhythmicity. Beyond clock times of stool collection, our study additionally included stooling history, dietary and sleep-related context, providing a comprehensive picture of diurnal dynamics within an age-controlled manner. Specifically this approach evidenced sleep history as strong driver of diurnal microbial dynamics, whereas the effect of prior fasting time was surprisingly negligible.

Our data reveal that the gut microbiota becomes more diverse and evenly distributed as the day progresses, indicated by increased bacterial diversity and evenness with later stool timings. This confirms diurnal dynamics in infant gut microbiota composition, as previously proposed based on data modelling [30]. Increased microbial diversity and evenness with later stool timings were proposed to be affected by feeding patterns [18], with morning defecation maintaining reduced diversity due to overnight fasting and lower microbial activity [13]. Yet, our novel results unravel that microbial effects of meal timing are primarily driven by age (details follow). Further, the link of gut microbiome composition with the stool timing is very clear with younger infants and disappear with infant maturation [28], suggesting that the early gut microbiome is particularly sensitive to timing dynamics [31].

The observed link between stool timing and zOTU abundance further points to emerging entero-diurnal (possibly circadian) rhythms. Lower microbial abundance in the morning is likely also intertwined with overnight fasting, with abundance increasing throughout the day as metabolic activity and food intake rise [12]. As the day progresses, increasing metabolic activity and nutrient intake change the local microbial environment, likely leading to higher zOTU counts later in the day. Overall, the effect of stool timing on microbial composition is a crucial discovery that could inform ongoing initiatives applying microbial composition as markers for infant health and disease. For example, gut microbiome can predict risks of gastrointestinal disorders like necrotizing enterocolitis [32], and knowledge of factors that fine-tune the variations in microbial diversity, like diurnal patterns, may improve precision in identifying infants at risk of dysbiosis [33]. It may also fundamentally supply the development of targeted prebiotics and probiotics for infants and young children, helping to optimize gut health, indirectly, developmental outcomes [20, 21].

We report a strong effect of sleep pressure on gut microbiome composition during infancy. Specifically, we highlight a microbial diversity increase with longer wake periods, indicating that microbial diversity benefits from resumed physiological processes during the waking period [34]. In addition, the results revealed that richness and evenness also rise after just a brief wakefulness duration. These findings highlight an effect of sleep-wake history on gut microbiome composition, which may connect to the gradual growth of microbial enteral clocks [35, 36]. We also detail insights into different age groups. Interestingly, the link of sleep pressure to diversity and richness at the age of 3 months, emphasizes a potentially critical role of sleep in early microbiome establishment. Sleep influences metabolic and immune functions, which in turn shape the gut microbiome [34, 37]. The reduced effect of sleep on microbial composition at older ages may reflect the microbiome’s stabilization and growing resilience as infants sleep patterns and the circadian regulator become more solid. In addition, the findings showed that zOTU abundance increases with sleep pressure, suggesting that the body’s metabolic state after waking supports this increase in abundance [38]. This post-waking surge in zOTUs could indicate that as metabolic processes resume, the gut environment becomes more suitable for microbial growth and diversity.

Finally, we determined the effect of meal history as a source of influence on gut microbiome composition. Time passed between meal and stool unraveled a positive correlation with gut microbial richness, suggesting that longer intervals between feeding promote a more diverse microbiome. This positive correlation reinforces the role of feeding in promoting microbial growth and diversity, highlighting how feeding patterns shape gut microbiota composition, as previously reported in [12, 27, 29, 39, 40] However, when the analysis was conducted separately for different age groups, this effect of meal history on microbial composition was no longer observed, indicating that age is a significant confounding factor in this relationship. One reason for this lack of effect could be that most samples collected for time since last meal fall within the first and second hour, which lead to a biased presentation of data. Unlike prior studies that predominantly examined feeding type (e.g., breastfeeding versus formula feeding [29]) or diet composition in later stages of adulthood [40], our study highlights the importance role of meal timing on early-life microbiota composition. The observations underscore the importance of considering feeding intervals alongside feeding type in infancy, as these patterns not only affect analyses but may also play a critical role in shaping early microbiota development. In addition, the results showed no significant association between zOTU abundance and prolonged fasting, except for a weak negative correlation, underscoring the reliance of microbial diversity on nutrient availability [41]. ZOTU abundance also declines as nutrient levels drop over time, suggesting fewer species can thrive in the less nutrient-depleted environment. In addition, our analysis revealed that temporal factors associate with microbiome composition at the phylum level, with age-specific patterns: lower stool frequency reduced *Proteobacteria* at 3 and 12 months, while *Actinobacteria* increased at 12 months, consistent with their early microbial succession. Sleep pressure reduced *Firmicutes* at 3 months, associating wake duration to microbiome composition. However, no consistent links were found for stool timing or time since last meal, suggesting that microbial responses to these factors may be more dynamic and age-dependent.

Overall, this study unravels the responsiveness of the infant gut microbiome to daily physiological and behavioral changes. The strongest effects were observed in the interplay between bowel movement frequency and microbial parameters, where longer intervals since the last stool were linked to increased microbial diversity and evenness. Stool timing (clock time) also had a strong impact on microbial composition, with diversity and evenness tending to increase later in the day. In addition, sleep pressure played a critical role, particularly at 3 months, where longer wake periods were associated with increased microbial richness and evenness. In comparison, meal history had a comparatively weaker effect on microbial richness, evident mostly in the immediate post-feeding period. Notably, the effects of bowel movement frequency, stool timing, and sleep pressure were more pronounced in younger infants, emphasizing the unique sensitivity of the early gut microbiome to temporal dynamic factors. These findings may inform future studies aimed at designing supplemental treatments to optimize early microbiome development and possibly long-term health.

## 5. Conclusion

The study highlights the association between stool dynamics and temporal factors that act as zeitgebers for establishing sleep rhythm across the first months of human life. Samples from 198 healthy infants ages 3-31 months provided the fundament to test associations between time since last stool, stool timing, sleep pressure, and time since last meal with gut microbiome composition (e.g., alpha diversity, richness, evenness; abundance and phyla). The study identified a negligible effect of immediate infant fasting history on microbial composition, but two strong microbial determinants in bowel frequency and sleep pressure. Notably, age-specific transitions emerged as the strongest predictors, with the sleep-gut microbiome link being most prominent—and thus most vulnerable—at the earliest ages. Our findings emphasize the heightened responsiveness of the infant gut microbiome to environmental cues, particularly at 3 months of age. In addition, the findings highlight the need for crucial adjustments when using microbiota profiles as indicators of health and development.

We call for future research to consider the role of temporal factors when conceptualising clinical applications related to infant microbial diagnostics and nutritional interventions. Longitudinal studies tracking the impact of temporal cues on the infant gut microbiome, with a focus on transitions to solid foods and regular sleep patterns, will be essential. In addition, investigating the establishment of circadian rhythms in diverse populations could provide a deeper understanding of microbiota-related health, further informing clinical practice and interventions. Finally, while this study focused on phylum-level bacterial composition, future studies should extend this approach to the genus level to capture more nuanced microbial shifts and their potential functional implications.

## Appendix

**Table 1.**
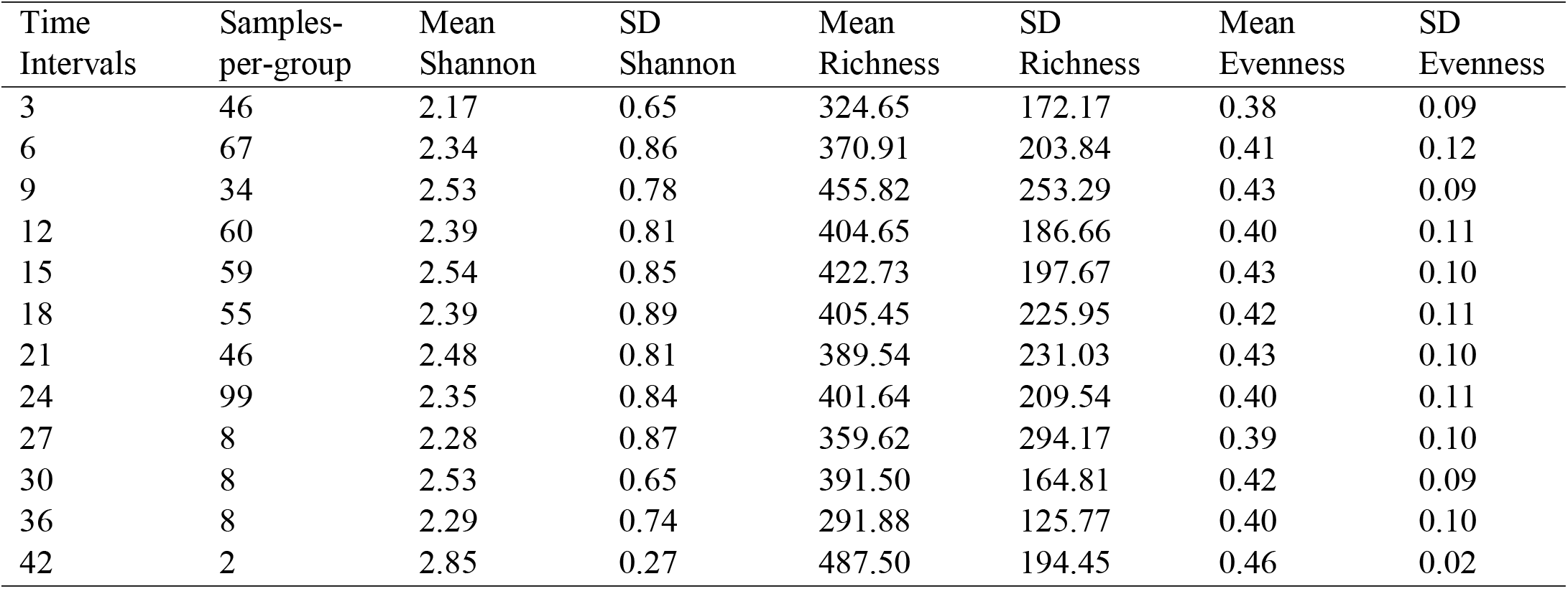
Summary statistics of microbial diversity, richness, and evenness across time intervals since last stool.

**Table 2.** Summary of microbial diversity, richness, and evenness across stool timing intervals.

| Time Intervals | Samples-per-group | Mean Shannon | SD Shannon | Mean Richness | SD Richness | Mean Evenness | SD Evenness |
| --- | --- | --- | --- | --- | --- | --- | --- |
| 3 | 2 | 1.32 | 0.43 | 172.5 | 16.26 | 0.26 | 0.08 |
| 4 | 4 | 1.69 | 0.49 | 288.25 | 167.64 | 0.34 | 0.11 |
| 5 | 1 | 1.98 | 0.51 | 204 | 170.8 | 0.37 | 0.10 |
| 6 | 5 | 2.47 | 0.89 | 441.6 | 374.55 | 0.41 | 0.09 |
| 7 | 32 | 2.39 | 0.68 | 422.25 | 158.48 | 0.41 | 0.1 |
| 8 | 61 | 2.48 | 0.9 | 415.13 | 217.56 | 0.43 | 0.11 |
| 9 | 76 | 2.53 | 0.86 | 437.33 | 214.22 | 0.43 | 0.11 |
| 10 | 44 | 2.2 | 0.91 | 360.41 | 219.3 | 0.39 | 0.12 |
| 11 | 22 | 2.26 | 0.71 | 344.32 | 206.76 | 0.4 | 0.1 |
| 12 | 44 | 2.2 | 0.75 | 324.14 | 179.85 | 0.4 | 0.1 |
| 13 | 49 | 2.43 | 0.86 | 382.35 | 214.52 | 0.42 | 0.13 |
| 14 | 34 | 2.54 | 0.82 | 426.53 | 214.05 | 0.43 | 0.1 |
| 15 | 22 | 2.21 | 0.83 | 361.32 | 161.11 | 0.43 | 0.1 |
| 16 | 29 | 2.33 | 0.85 | 364.83 | 246.46 | 0.41 | 0.11 |
| 17 | 30 | 2.5 | 0.67 | 380.07 | 144.71 | 0.43 | 0.09 |
| 18 | 28 | 2.22 | 0.77 | 337.29 | 207.78 | 0.39 | 0.11 |
| 19 | 32 | 2.39 | 0.67 | 406.69 | 199.66 | 0.4 | 0.09 |
| 20 | 26 | 2.49 | 0.81 | 443.81 | 240.15 | 0.42 | 0.1 |
| 21 | 17 | 2.01 | 0.76 | 303.06 | 179.65 | 0.35 | 0.11 |
| 22 | 1 | 3 | 0.77 | 266 | 180.2 | 0.54 | 0.12 |
| 23 | 5 | 2.06 | 0.61 | 303.6 | 116.26 | 0.36 | 0.09 |

